# Intrahost mutational dynamics parallel long-term genome evolution in endosymbionts

**DOI:** 10.64898/2026.01.27.701382

**Authors:** Younghwan Kwak, Gordon Bennett

**Affiliations:** Department of Life and Environmental Sciences, University of California, Merced, CA, USA

## Abstract

Obligate endosymbionts of insects undergo extreme genome evolution, marked by accelerated molecular evolution, severe base-pair compositional bias, and massive gene loss. However, the microevolutionary processes driving these patterns remain poorly understood, as they occur at intrahost population scales that are rarely captured. To address this gap, we measured intrahost genetic diversity of two endosymbionts, *Karelsulcia* and *Nasuia*, from the aster leafhopper, *Macrosteles quadrilineatus* (Hemiptera: Cicadellidae). Contrary to the theoretical expectation of strict clonality, we found that both endosymbionts harbor measurable intrahost genetic variation, with lineage-specific mutational dynamics that parallels long-term evolutionary trends. *Karelsulcia* showed sparse intrahost variation dominated by repeat-associated indels, while *Nasuia* exhibited more abundant single-nucleotide mutations that appear to shape genome-wide A+T bias. Mitochondrial heteroplasmy did not covary with endosymbiont nucleotide diversity, indicating that these patterns are not driven by host-level dynamics. Notably, recurrent nonsynonymous variants in *Nasuia* affect essential genes for amino acid biosynthesis and translation. The intrahost mutational patterns we observed in endosymbionts are consistent with long-term sequence changes between our population founder genome and contemporary endosymbiont populations after ∼11 years of maintenance. Taken together, our results demonstrate how distinct mutational processes operating at the intrahost populations scale drive macroevolutionary patterns in endosymbiont genomes. Moreover, our study establishes that laboratory endosymbiont systems provide a powerful framework for dissecting and understanding these fundamental evolutionary processes.

## Introduction

Obligate intracellular symbionts (endosymbionts) in insects represent some of the most striking cases of genome evolution in bacteria. Due to their host restriction over millions of years, endosymbionts evolve under extreme demographic constraints that lead to strong genetic drift: strict vertical transmission, repeated population bottlenecks, and minimal opportunities for recombination (Baumann 2005; Moran et al. 2008). Comparative genomics has yielded a detailed picture of the long-term consequences of these conditions. Across insects, endosymbiont genomes are stripped of up to 90% of their genetic and functional capabilities compared with their free-living relatives (reviewed by McCutcheon et al. 2024).

They further show general patterns of accelerated molecular evolution, pronounced deletion bias, and strong A+T nucleotide compositional bias (Moran 1996; Mira et al. 2001; Kuo and Ochman 2009). The processes that drive these evolutionary outcomes can be complex and distinct between endosymbiont lineages of different hosts and origins. For example, complementary endosymbionts that share the same host can diverge sharply in evolutionary rate and functional capacity, indicating that genome erosion is neither uniform nor fully predictable across lineages (e.g., Sloan and Moran 2012; Nakabachi et al. 2013; Bennett et al. 2014; Szabó et al. 2022). These macroevolutionary patterns underpin our current view of how endosymbiosis shapes bacterial genomes and how mutation, drift, and selection interact under chronic host dependence (Wernegreen 2015; Wilson and Duncan 2015).

Despite a dramatic proliferation in endosymbiont genomes and the expanding theories they have yielded (*e.g.,* see Wernegreen 2002; McCutcheon et al. 2019; Kaltenpoth et al. 2025), we have little direct evidence for the microevolutionary processes that generate them. This limitation arises primarily from a mismatch between the scale at which evolution operates (i.e., population-level) and the more often measured outcomes that it produces. The mutations that ultimately lead to endosymbiont evolution (e.g., selection to maintain function and drift-related gene loss; Sabater-Muñoz et al. 2017) arise and segregate within individual hosts. However, the majority of endosymbiont genomic studies have relied on pooled individuals and summarize variation as consensus genomes, yielding composite representations of only the dominant variants within host populations (Moran et al. 2009; Williams and Wernegreen 2012; Williams and Wernegreen 2013). As a result, it remains empirically unmeasured in nearly all endosymbiotic systems how frequently endosymbionts acquire variation, how that variation is distributed across loci, and the evolutionary fate of that variation at the intrahost population scale (but see Woyke et al. 2010; Perreau et al. 2021). Since individual hosts ultimately control endosymbiont function, reproduction, and fitness (*e.g.,* Vigneron et al. 2014; Smith and Moran 2020; Rafiqi et al. 2022; Oishi et al. 2023), this level of variation is critical to understanding how evolutionary processes shape endosymbiont genomes.

Here, we address this limitation by leveraging a tractable laboratory system well suited for direct measurement of endosymbiont genetic variation at the intrahost scale. The aster leafhopper *Macrosteles quadrilineatus* harbors two obligate bacterial endosymbionts, *Candidatus* Karelsulcia muelleri (Bacteroidetes; hereafter referred to as *Karelsulcia*) and *Candidatus* Nasuia deltocephalinicola (Betaproteobacteria; hereafter *Nasuia*), each with extremely reduced genomes (190 kb and 112 kb, respectively). Similar to most other endosymbionts in the Auchenorrhyncha suborder, *Karelsulcia* and *Nasuia* complement each other to provide their hosts with the 10 essential amino acids that are lacking in their xylem and phloem plant sap diets (Bennett and Moran 2013). Both genomes maintain a core set of essential nutritional genes, but are lacking genes in most other essential functions that include translation and transcription, energy synthesis, and DNA replication and repair (Bennett and Moran 2013). By quantifying and characterizing how these degenerate genomes harbor and partition genetic variation within individual hosts, we reveal intrahost mutational dynamics that underpin macroevolutionary patterns of genome evolution in *Karelsulcia* and *Nasuia*. Our results demonstrate that laboratory endosymbiont systems can serve as uniquely powerful frameworks for studying how microevolutionary processes shape macroevolutionary outcomes at the genome-wide evolution scale.

## Results

### Establishment of a framework for quantifying intrahost endosymbiont variation

Understanding how obligate intracellular symbionts carry and segregate genome-wide variation requires an experimental design that can separate true endosymbiont variation from host-level genetic heterogeneity (i.e., genetic structure within host populations) and sequencing artifacts. To accomplish this aim, we developed a high-resolution framework that integrates three key features below.

First, we minimized host demographic noise by analyzing a cross-sectional sample from a single laboratory lineage (20 adult individuals; 10 females and 10 males). Our lab colony has been isolated from wild populations for ∼11 years (Bennett and Moran 2013). This bottlenecked host background constrains inherited host genetic diversity and limits cryptic population structure introduced by host demographics, so differences among individuals are more likely to reflect mutations that arose during isolation rather than divergent host genotypes as would be expected in natural populations. At the same time, the colony’s small effective population size can amplify stochastic mutation accumulation over laboratory generations, making new variants more likely to drift to detectable frequencies.

Second, we generated population consensus genomes from our study cohort for read mapping and variant calling. Population consensus genomes are widely used in human genetic studies to reduce bias caused by mismatches between a universal reference and the focal population (Cho et al. 2016; Levy-Sakin et al. 2019; Bergström et al. 2020; Takayama et al. 2021; Kaminow et al. 2022). We applied the same rationale here because the previously available public NCBI genomes of endosymbionts and mitochondria were assembled from the wild founder population (Bennett and Moran 2013). By building *de novo* consensus genomes from our dataset, we ensured that our analyses are anchored to the contemporary lab lineage.

Third, we compared genetic variation patterns between the two endosymbionts, *Karelsulcia* and *Nasuia*, as an internal contrast. *Nasuia* has higher AT-bias and a greatly accelerated rate of molecular evolution compared with *Karelsulcia,* which is drastically more stable (Bennett et al. 2014; Vasquez and Bennett 2022). This contrast allowed us to determine whether genetic variation reflects endosymbiont-intrinsic processes or shared environmental influence (*e.g.,* bacteriocyte metabolic environment). We further analyzed intrahost genetic variation in the host mitochondrial genome (i.e., mtDNA heteroplasmy) as an additional internal benchmark for both endosymbionts (Gray 2012; Raval et al. 2023).

We verified that our framework rests on robust genomic and technical foundations. We performed deep metagenomic sequencing in duplicate. Median per-base coverage (IQR) across two sequencing runs was 1,496× (946–2,749×) for *Karelsulcia*, 472× (344–1,102×) for *Nasuia*, and 7,735× (3,785–10,558×) for mtDNA (**Supplementary Table 1; Supplementary Fig 1**). Our *de novo* population consensus genomes for *Karelsulcia, Nasuia*, and mitochondria showed > 99% pairwise identity to their respective founder references (**Supplementary Table 2**). We retained only variants independently recovered across the two sequencing runs under identical filtering, yielding a conservative, reproducible callset (**Supplementary Table 3;** Roder et al. 2023). Variant count per kb (i.e., variant density) did not correlate with sequencing depth (Spearman ρ = 0.27, 0.28 and-0.28 for *Nasuia*, *Karelsulcia*, and mitochondria, respectively, all P > 0.23). Finally, multivariate QC of variant presence–absence profiles indicated no systemic effects of library batch, sample cohort, or host sex on the structure of variant profiles (**Supplementary Fig. 2**; **Supplementary Table 4**).

### Contrasting mutational landscapes of *Karelsulcia* and *Nasuia*

Contrary to the theoretical expectation of strict clonality, both bacterial symbionts harbored measurable numbers of variants within the same host individual (**Fig. 1**). In 15 of the 20 populations, we found higher total variant density in *Nasuia* than *Karelsulcia* (Wilcoxon signed-rank, P = 0.006; median ratio 1.46×; **Fig. 1A**). We next examined the composition of these variants. We classified observed variants into three classes: single-nucleotide variants (SNVs), multi-nucleotide variants (MNVs; two or more adjacent SNVs), and small insertions and deletions (indels; < 50 bp). Large structural variants (*i.e.,* indels ≥ 50 bp; Alkan et al. 2011) were outside of the scope of this study and were not estimated. We found that *Nasuia* was relatively SNV-enriched, whereas *Karelsulcia* was indel-enriched (SNV fractions: medians 0.46 [IQR 0.40-0.51] vs 0.20 [0.15-0.29]; indel fractions: 0.51 [0.42-0.54] vs 0.77 [0.69-0.82]; Wilcoxon, P < 10^-4^ for both; **Fig. 1B**). Multi-nucleotide variants (MNVs) were rare and comparable between endosymbionts (median fraction ∼0.05; Wilcoxon, P = 0.78; **Fig. 1B**).

**Figure 1.**
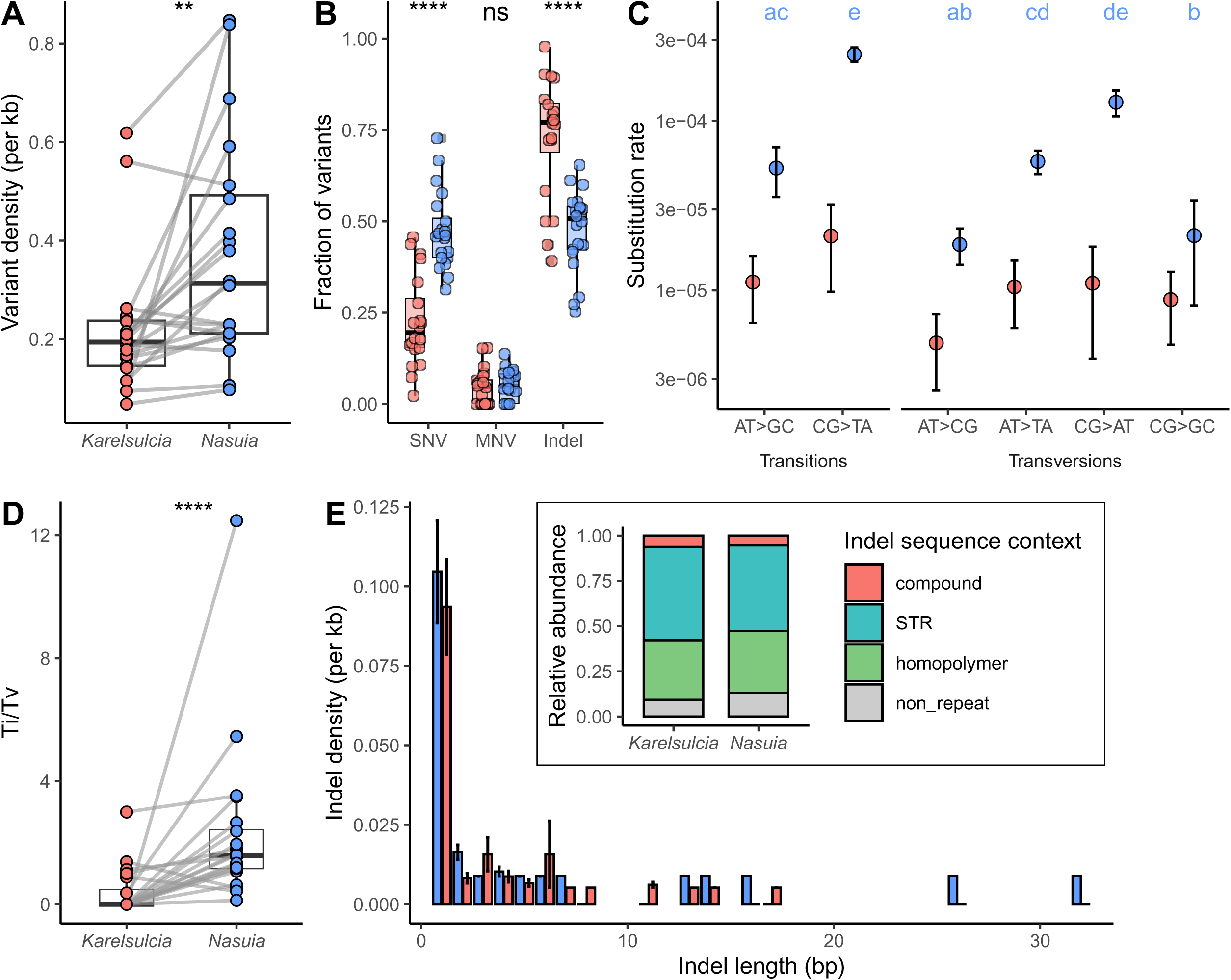
Intrahost genetic variation in *Karelsulcia*, and *Nasuia.* **(A)** Variant density across 20 endosymbiont populations. Total number of variants were normalized to the respective genome length and scaled to variants per kilobase. Each line connects paired *Karelsulcia* and *Nasuia* populations from the same host (**, *P* < 0.01 Wilcoxon singed-rank test). **(B)** Composition of variant classes within each endosymbiont. Fractions of single-nucleotide variants (SNVs), multi-nucleotide variants (MNVs), and insertions and deletions (indels) were calculated relative to total variants per population (ns, *P* ≥ 0.05; ****, *P* < 0.0001; Wilcoxon signed-rank tests). **(C)** Six directional SNV spectra. SNV rates were normalized by the number of possible sites for each nucleotide type (A/T or C/G) within each genome. Letters indicate statistically distinct groups (Kruskal–Wallis followed by Dunn’s post hoc test, *P* < 0.05) **(D)** Transition/transversion (Ti/Tv) ratio across hosts. Each line connects paired endosymbionts (****, *P* < 0.0001; Wilcoxon signed-rank test). **(E)** Indel length distribution. Indel frequencies were normalized to total genome to the respective genome length and scaled to variants per kilobase. Stacked bar indicates the relative contribution of homopolymer, short tandem repeat (STR), compound repeat, and non-repeat contexts.

We then evaluated base-level mutation biases using the six directional SNV classes (A>C, A>G, A>T, C>A, C>G, and C>T, after collapsing reverse complements). *Nasuia* and *Karelsulcia* differed in their overall SNV spectra (χ^2^ = 11.0, df = 5, P = 0.039; **Supplementary Table 5**). This difference was spread across all base-change classes rather than driven by any single type (Benjamini-Hochberg q > 0.05 after Fisher’s exact test). Within *Nasuia*, SNV rates varied strongly among base-change classes, driven by elevated C>T and C>A changes (Dunn post-hoc FDR < 0.01 after Kruskal-Wallis; **Fig. 1C).** In contrast, *Karelsulcia* showed no significant variation across base-change classes (Kruskal-Wallis, χ^2^ = 3.50, df = 5, P = 0.62). The transition/transversion (Ti/Tv) ratio was consistently higher in *Nasuia* than in *Karelsulcia* across hosts (18 out of 20 endosymbiont pairs; Wilcoxon, P = 4.10×10^-5^; median paired ΔTi/Tv = 1.45 [IQR 0.60-2.06]; **Fig. 1D**).

Observed insertion-deletion (indels) were mainly single-base events in both bacterial symbionts, with rare longer indels observed only in *Nasuia* (up to 32 bp; **Fig. 1E).** Normalized indel-length distributions by genome size did not differ significantly between *Karelsulcia* and *Nasuia* (Fisher’s exact P > 0.05). Homopolymer tracts and short tandem repeats (STRs) are known hotspots for indels because replication slippage is common in low-complexity DNA (Redelings et al. 2024). To quantify this effect, we normalized indel counts by the total length of each repeat class in the population consensus genomes (*i.e.,* the summed length of homopolymers or STRs with a track length ≥ 4 bp). These normalized profiles show that repeat-associated indels account for ∼90% of the total indel density in both *Nasuia* and *Karelsulcia* (**Fig. 1E**, see inset).

We further characterized mitochondrial variation to contextualize patterns observed in endosymbionts (**Supplementary Fig. 3A**). Because the mitochondrial variant detection pipeline does not permit MNV detection, here we split *Nasuia* and *Karelsulcia* MNVs into their constituent SNVs to generate a comparable two-class profile of SNVs and indels. Median fractions were [SNV = 0.472, indels = 0.528] for mitochondria, [SNVs = 0.530, indels = 0.470] for *Nasuia*, and [SNV 0.275, indels = 0.725] for *Karelsulcia*. In 15 out of 20 hosts, mitochondria were compositionally closer to *Nasuia* than to *Karelsulcia* (paired L1 distance on [SNVs, indels]; Wilcoxon signed-rank, P = 0.01; **Supplementary Fig. 3B-C**).

### Frequency and spatial heterogeneity of intrahost variation

Our catalog of intrahost variants shows that bacterial symbiont populations within individuals are not genetically inert. To determine the prevalence of each variant within its respective population, we examined the allele frequency (AF) spectrum. *Nasuia* exhibited a broader AF distribution in contrast to *Karelsulcia*, which was almost entirely skewed toward rare alleles (**Fig. 2A**). Intermediate-frequency variants (AF = 0.25-0.75) were more common in *Nasuia*, appearing at higher levels in 19 of 20 populations (Wilcoxon, P = 5.6 × 10^-5^; median Δ 8.78 [95% CI 6.54-Inf]; **Fig. 2B**). *Karelsulcia* carried no intermediate-frequency variants in 18 populations (**Fig. 2B**). The mitochondrial AF spectrum resembled that of *Karelsulcia*, dominated by low-frequency heteroplasmic variants with very few intermediates (**Fig. 2A-B**). Among *Nasuia*, intermediate-allele fractions did not differ by sex (Wilcoxon, P > 0.1), indicating population-specific dynamics rather than host sex explain most of the observed variation.

**Figure 2.**
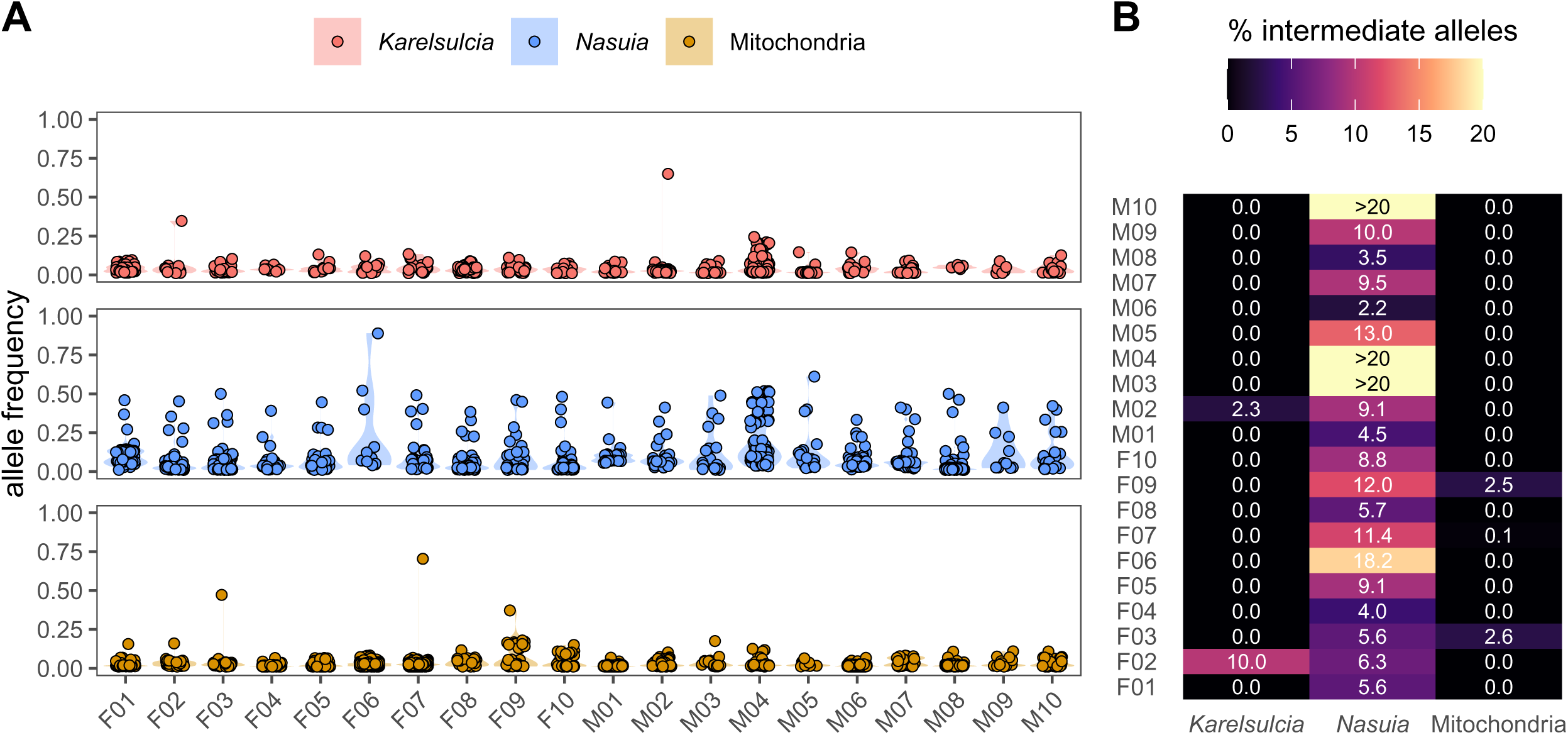
Allele frequency distributions for *Karelsulcia, Nasuia*, and mitochondria. **(A)** Allele frequency spectra across 20 populations. Each point represents a segregating site in an individual host. The y-axis indicates the frequency of the alternate allele. **(B)** Fraction of intermediate-frequency alleles (0.25 < AF < 0.75) per population. Values represents the percentage of intermediate-frequency variants relative to all detected variants in each species.

We next quantified the overall magnitude of intrahost variation using nucleotide diversity (π).

Across all 20 populations, *Nasuia* showed a 6.3-fold higher genome-scaled π than *Karelsulcia* (Wilcoxon, P = 4.8×10^-5^; median paired Δπ=8.9×10^-5^ [1.4×10^-5^-Inf]; **Fig. 3A**). Likewise, *Nasuia* harbored a greater proportion of segregating 1 kb windows (π > 0; 20 out of 20 pairs; Wilcoxon, P = 4.8×10^-5^; median Δ = 0.11 [0.094-Inf]; **Fig. 3B**), indicating that variation spans a larger fraction of its genome. We observed a weak trend toward positive spatial clustering of π windows in *Nasuia*, but not in *Karelsulcia* (Moran’s I = 0.10 vs-0.02, both P > 0.05). Although this effect was not statistically significant, the positive clustering indicates that *Nasuia*’s segregating windows tend to cluster in its genome. Additionally, *Nasuia* consistently shared many of the same segregating windows than did *Karelsulcia* (mean Jaccard similarity = 0.27 vs 0.11; Wilcoxon, P = 1.3 ×10^-42^; median ΔJ = 0.169), indicating recurrent mutation at common genomic positions. *Nasuia* showed more uneven distribution of π across segregating windows than did *Karelsulcia* (Gini = 0.69 vs 0.53; Pielou’s J’ = 0.40 vs 0.62; **Figure 3C-D**), with a subset of regions carrying disproportionately high diversity.

**Figure 3.**
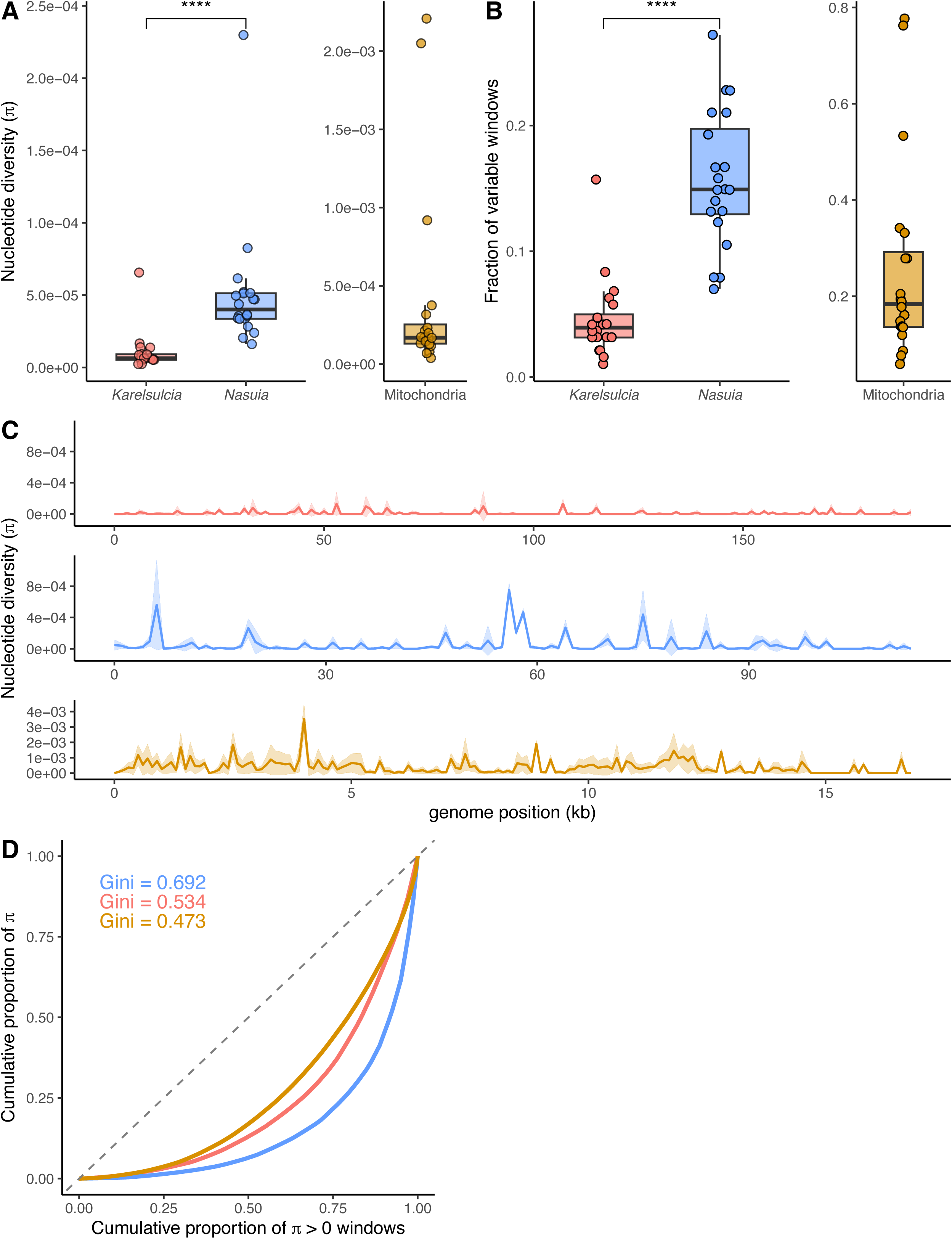
Nucleotide diversity (π) in *Karelsulcia*, *Nasuia*, and mitochondria. (A) Genome-wide π across 20 populations. **(B)** Fraction of variable 1 kb windows (π > 0) per genome (****, *P* < 0.0001; Wilcoxon signed-rank test). Statistical comparisons in panels **(A)** and **(B)** were performed only between *Karelsulcia* and *Nasuia*; mitochondrial values are shown with a scale offset for visualization. **(C)** Distribution of π across genomic position in *Karelsulcia* (top) and *Nasuia* (middle), and mitochondria (bottom). Lines and shaded regions represent mean ± s.e. across populations. **(D)** Lorenz curves showing the cumulative distribution of π across 1 kb windows. Gini coefficients values are annotated.

Due to its elevated mutational load, the mitochondrial genome harbored higher overall nucleotide diversity than either endosymbiont (genome-wide pi = 1.7×10^-4^; **Fig. 3A**). However, mitochondrial diversity did not covary with that of either endosymbiont across hosts (Spearman ρ =-0.06 and 0.13 for *Nasuia* and *Karelsulcia*, respectively, both P > 0.5). This result indicates that host-to-host variation in mitochondrial heteroplasmy is independent of endosymbiont diversity levels. Across segregating windows, mitochondrial heteroplasmy was relatively uniform in magnitude (Gini = 0.47; Pielou’s J’: 0.67), but showed modest localized diversity rather than being randomly scattered (Moran’s I = 0.17, P = 0.01).

Together, these results suggest that while mitochondrial genomes accumulate more mutations overall, their per-window diversity is more evenly distributed than in the endosymbionts, even as segregating windows cluster along the genome.

### Functional bias of mutations in *Nasuia*

The presence of local diversity across endosymbiont genomes raises the question of whether these mutation hotspots reflect neutral turnover or affect protein-coding function. To address this question, we compared AF-weighted rates of synonymous (pS) and nonsynonymous (pN) SNVs across all coding regions using a codon-aware framework (see **Materials and Methods**). Across both endosymbionts, variants with high pS were confined to a small number of genes and typically appeared as private SNVs in only a few endosymbiont populations. Per-population pS, which averages across all genes within each population, did not differ significantly between endosymbionts (Wilcoxon, P = 0.50). In contrast, nonsynonymous variants were broadly recurrent across genes but exhibited uniformly low pN (**Fig. 4A**). Although pN values were low in both endosymbionts, per-population pN was significantly higher in *Nasuia* than in *Karelsulcia* (Wilcoxon, P < 0.0005). Genes that carried nonsynonymous SNVs showed little overlap with those carrying synonymous SNVs, indicating that functional and neutral turnover tend to affect different loci at the intrahost scale.

**Figure 4.**
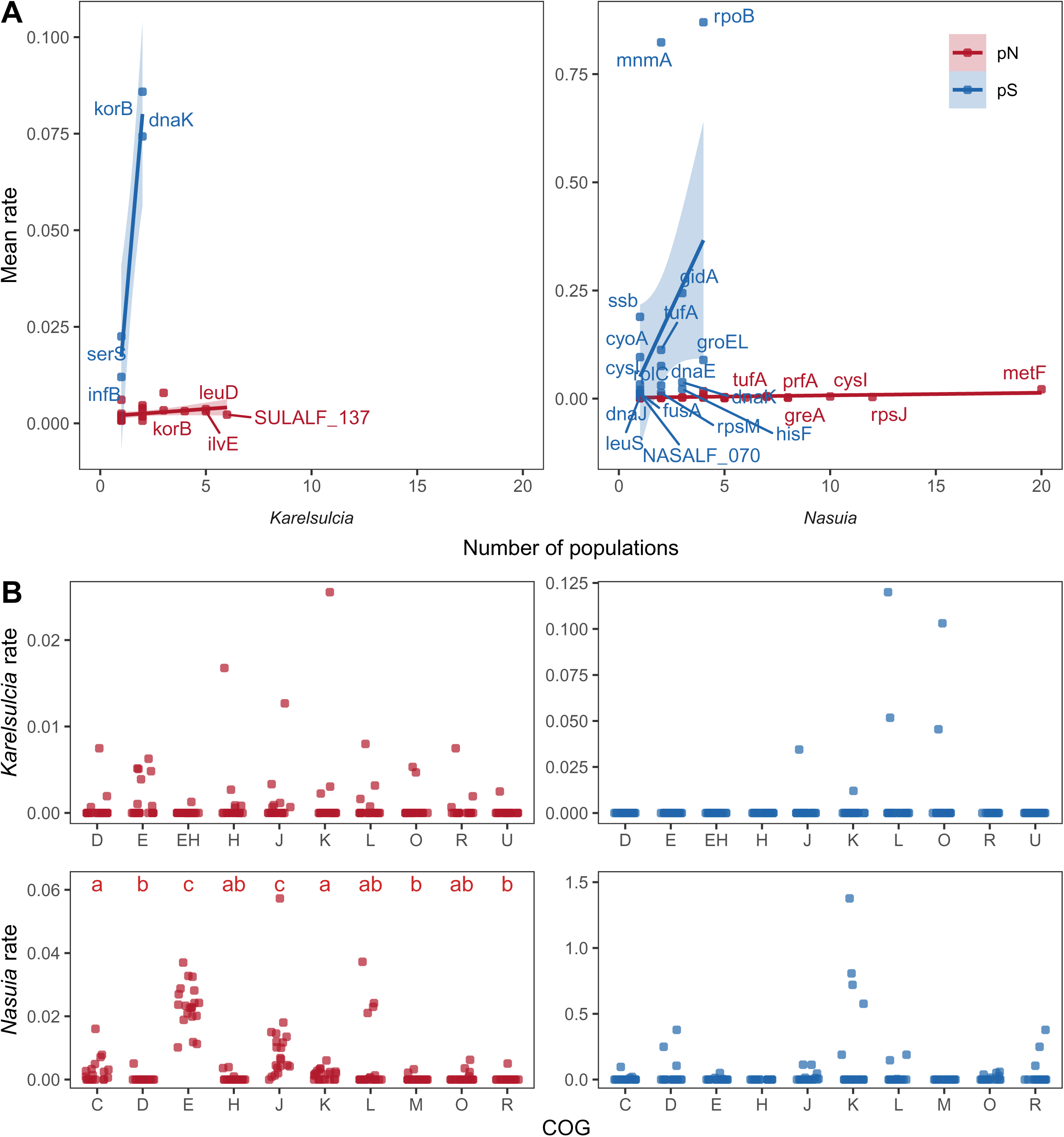
Coding-sequence variation in *Karelsulcia* and *Nasuia*. **(A)** Gene-level nonsynonymous (pN, red) and synonymous (pS, blue) rates plotted against the number of populations with detected variants. Lines represent linear regression fits, and shaded regions show the 95% confidence interval. **(B)** Distribution of pN (left) and pS (right) across functional categories (COG): C, Energy production and conversion; D, Cell cycle control, cell division, chromosome partitioning; E, Amino acid transport and metabolism; H, Coenzyme transport and metabolism; J, Translation, ribosomal structure and biogenesis; K, Transcription; L, Replication, recombination and repair; M, Cell wall/membrane/envelope biogenesis; O, Posttranslational modification, protein turnover, chaperones; R, General function prediction only; U, Intracellular trafficking, secretion and vesicular transport). Letters indicate statistically distinct groups within *Nasuia* (Kruskal–Wallis followed by Dunn’s post hoc test, *P* < 0.05).

Several *Nasuia* genes accumulated recurrent nonsynonymous SNVs in multiple populations. The most extreme case was *metF*, which carried nonsynonymous variants in all 20 populations (**Fig. 4A**). This gene is located near a known repeat region, which likely elevates its mutability. Several informational genes (*tufA*, *rpsJ*, and *prfA*) and the sulfur-metabolism gene *cysI* also had recurring SNVs in ∼30–60% of hosts. Consistent with these patterns, nonsynonymous variations in *Nasuia* were concentrated in the Clusters of Orthologous Groups (COG) associated with amino acid metabolism (E) and Translation (J; Kruskal–Wallis, Dunn’s post hoc, P < 0.05; **Fig. 4B**). These categories encompass pathways that underpin *Nasuia*’s core nutritional role in the symbiosis (E) and that are among the most frequently reduced in streamlined bacterial genomes (J). In *Karelsulcia*, pN values showed no significant differences among COG categories.

We next tested whether indels in protein-coding regions have a functionally biased impact on genes and cellular functions. Gene-level coding indel rates did not differ among COG categories in either endosymbiont (Kruskal–Wallis, P > 0.1; **Supplementary Fig. 4**). Likewise, the fraction of genes with coding indels relative to all genes in each COG was not significantly enriched in any category in either endosymbiont (Fisher’s exact test, all FDR > 0.1). The diffuse distribution of coding indels across COG categories in both bacteria suggests that most indels are governed by local sequence architecture and structural tolerability rather than pathway-specific differences in selection.

### Temporal signatures of mutation fixation

Taking advantage of a reference genome from the original laboratory founder population, we asked how much of the contemporary genetic variation in *Karelsulcia* and *Nasuia* reflects mutations that emerged during ∼11 years of laboratory maintenance. Given their distinct inheritance and evolutionary dynamics, mitochondrial genomes were excluded from this temporal analysis. Using the founder reference as a baseline, we identified sites where contemporary alleles now represent the major allele (AF > 0.75; **Fig. 5**).

**Figure 5.**
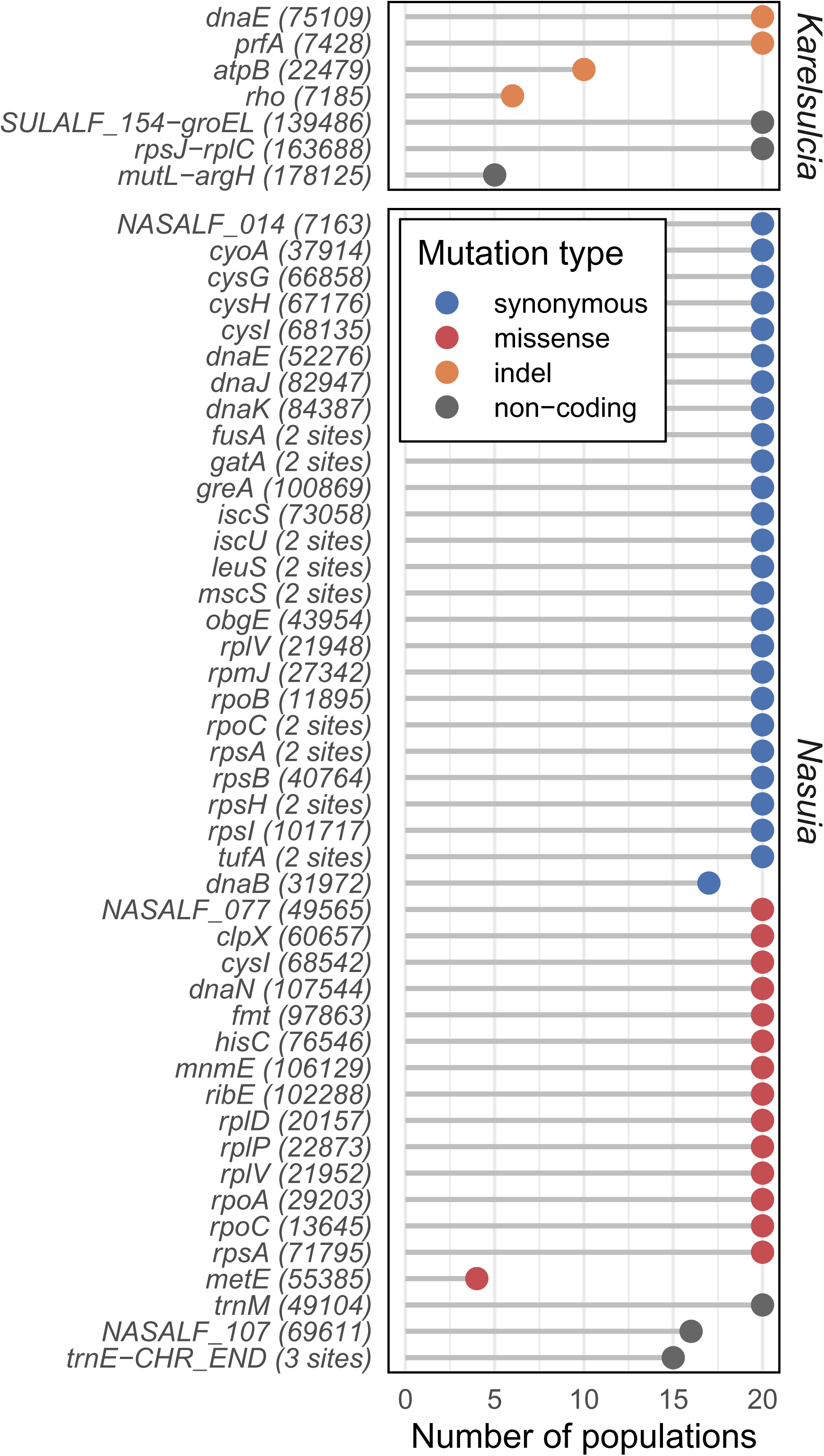
Founder–contemporary substitutions across *Karelsulcia* and *Nasuia*. Dot plots show the number of endosymbiont populations in which each nearly fixed founder–contemporary variant occurs. Points are colored by mutation type. Each row corresponds to a gene, with the genomic position of the variant shown in parentheses. Genes with multiple founder-contemporary sites are labeled with the number of sites. Founder–contemporary variants were identified by comparing contemporary endosymbiont populations to the laboratory founder reference sequences for *Karelsulcia* (GenBank: CP006060.1) and *Nasuia* (GenBank: CP006059.1).

In *Karelsulcia,* we detected only seven founder-contemporary differences, all of which are indels. These sites are distributed across four genes and three intergenic regions, including loci encoding core housekeeping functions such as DNA replication (*dnaE*), translation termination (*prfA*), ATP synthesis (*atpB*), and transcription termination (*rho*). The ATP synthase subunit gene *atpB* is notable because it carries two distinct alternative alleles relative to the founder sequence. Both involve expansion of a 6-bp tandem repeat (ATTACT): the founder allele contains six repeat units, whereas contemporary alleles carry either seven (+1) or nine (+3) repeat units (**Supplementary Table 6**). Together, these alleles occur in 10 populations, with one allele reaching AF = 0.80 in two populations and the other reaching AF = 0.94 in eight populations, suggesting that both are currently sweeping toward fixation.

As expected, *Nasuia* showed substantially greater accumulated genetic variation than *Karelsulcia*.

We identified 56 sites at which the contemporary allele is nearly fixed (AF = ∼0.99), 51 of which are present in all 20 populations (**Supplementary Table 6**). These founder-contemporary differences are dominated by single-nucleotide substitutions: 35 synonymous,15 missense, and three noncoding substitutions in intergenic or untranslated regions (UTRs). Only two intergenic indels were observed near the chromosome junction. Most genes carry a single synonymous substitution, but 11 genes showed multiple changes. As in our AF-weighted pN and pS analyses, genes with missense founder–contemporary changes are largely distinct from those with only synonymous changes, with overlap restricted to these four loci (*cysI*, *rplV*, *rpoC*, and *rpsA*). Missense founder–contemporary variants occurred primarily in genes assigned to COG J (translation; six genes) and E (amino-acid metabolism; three genes), the same functional categories that carry segregating nonsynonymous variants in the contemporary populations (**Fig. 4B**). This overlap suggests recurrent mutational input at these loci across both longitudinal and contemporary sampling intervals. Together, these results provide empirical evidence that short-term mutational processes in *Nasuia*—and the relative absence of such processes in *Karelsulcia*—mirror the contrasting long-term evolutionary dynamics of these symbionts.

## Discussion

Genomic investigation of bacterial endosymbionts in insects has detailed the macroevolutionary impacts of living in close host-association for millions of years. But the microevolutionary processes that derive these outcomes occur within and between individual hosts and have remained obscured. Within hosts, strictly vertically transmitted endosymbionts are often considered to be nearly clonal populations and, in theory, likely to inherit limited standing genetic variation (Woyke et al. 2010). However, endosymbionts are widely known to have elevated mutation rates that yield continual input of variation (Moran 1996). These processes can lead to detectable intra-and inter-host genetic variation with significant evolutionary consequences. Using deep metagenomic sequencing, we show that endosymbionts retain non-trivial, reproducible intrahost genetic variation. *Nasuia,* in particular, exhibited continuous single-base mutational input that is heavily A+T-biased. Even *Karelsulcia*, previously characterized by exceptionally low rates of molecular evolution (Bennett et al. 2014; Vasquez and Bennett 2022), carried detectable indel-biased intrahost mutations. These variants occurred in a non-random, recurrent manner and frequently affect core metabolic and informational functions. Some of these changes have swept toward fixation over ∼11 years of laboratory maintenance. Thus, intrahost diversity emerges as a dynamic layer of variation that parallels and shapes long-term endosymbiont genome evolution.

Detection of genetic variation in insect endosymbionts is constrained by both biological and technical limitations. Low bacterial symbiont DNA input from a single, small insect restricts the number of unique molecules sampled relative to the more abundant host nuclear and mitochondrial genomes. Achieving adequate evidence per site, therefore, requires ultra-deep sequencing. Even with modern, low-error sequencing platforms, pushing depth under low-input conditions inevitably amplifies risks of library-preparation and sequencing biases, including PCR amplification, duplication inflation, and allelic dropout (Robasky et al. 2014). In our study system, *Karelsulcia* is consistently more abundant than *Nasuia* (**Supplementary Fig. 1)**. Thus, the power to detect rare variants between endosymbionts is inherently asymmetric. Attempts to normalize samples by forcing equal depth has the potential to erase genuine biological differences rather than control for technical artifacts. Moreover, the biological properties of endosymbiont genomes add further challenges. They are extremely AT-rich and contain many low-complexity tracts (Bennett and Moran 2013). In such contexts, short reads map less reliably and homopolymer-associated errors increase (Stoler and Nekrutenko 2021), requiring additional methodological steps to accurately estimate genetic variation.

Here, we established an empirical framework to conservatively quantify intrahost genetic diversity in endosymbionts that overcomes biological and technical constraints. Most studies of endosymbiont mutational landscapes have focused on broad evolutionary scales, such as interspecific divergence or comparisons among endosymbiont strains from distinct host populations (Moran et al. 2009; Williams and Wernegreen 2012; Williams and Wernegreen 2013; Perreau et al. 2021). To our knowledge, only two previous analyses operate at a comparable scale to ours. First, a single-cell genomic assembly of *Karelsulcia* in the green sharpshooter showed no SNVs relative to the same single-host bacteriome metagenomic sequence data (Woyke et al. 2010). The authors interpreted this as evidence that the intrahost population is nearly clonal. However, this analysis relied on ∼20x coverage and was powered to detect variants with AF ≥ 0.25, based on detection simulation from two *E. coli* strains. In our dataset, most *Karelsulcia* variants fall below this threshold and are only resolvable at the ∼100-1,000x depths. Second, duplex sequencing of pooled bacteriomes from *Macrosteles sp. nr. severini* likewise detected rare *de novo* mutations in *Karelsulcia* and *Nasuia* (Waneka et al. 2021). The resulting SNV and indel profiles closely match our intrahost patterns, including the stronger A+T-biased SNV pattern in *Nasuia* and repeat-associated indels in both partners (**Fig. 1**). Together, the duplex-seq and our ultra-deep intrahost data indicate that this mutational asymmetry is a consistent feature of the *Karelsulcia*–*Nasuia* symbiosis across host lineages, sampling designs, and analytical scales.

*Karelsulcia*’s mutational input is strongly suppressed, and variation instead appears to be dominated by rare indel events. Its apparent indel bias may be best understood as a consequence of genome architecture. *Karelsulcia* has a larger genome with more extensive homopolymer and STR tracts. There is, therefore, more physical substrate on which replication slippage can occur. After scaling indel counts by the total length of these repeat tracts, *Karelsulcia* and *Nasuia* show comparable repeat-normalized indel rates (**Supplementary Fig. 4**). This pattern indicates that the excess number of *Karelsulcia* indels reflects its repeat substrate, not an elevated intrinsic propensity for indel formation (e.g., loss of the DNA mismatch repair [MMR] system in many obligate insect endosymbionts; reviewed in McCutcheon and Moran 2012). Against this background, the depressed rate of coding SNV turnover suggests that most point mutations in *Karelsulcia* are either strongly suppressed or efficiently removed.

Thus, *Karelsulcia*’s streamlined biosynthetic repertoire evolves primarily through occasional replication slippage in repetitive regions rather than continual amino-acid replacement.

In contrast, *Nasuia*’s greater intrahost genetic diversity relative to *Karelsulcia* is consistent with an elevated mutation rate, potentially due to reduced DNA repair capacity. Synonymous substitution rates (dS or Ks) are commonly used as a proxy for underlying mutation rates, particularly under the assumption that synonymous changes are largely neutral (Hershberg and Petrov 2008). Although the distribution of gene-level pS values did not differ significantly between endosymbionts (Wilcoxon test), *Nasuia* exhibited a greater number of genes with detectable synonymous variation and a substantially broader pS range when variation was present (**Fig. 4A**). Moreover, in lineages with extremely small effective population sizes (*N_e_*), the high mutational rates strongly interact with drift and could expose core pathways to continual turnover (Moran 1996). In our system, the ∼1:3 sequencing-depth ratio between *Nasuia* and *Karelsulcia* may reflect a smaller standing population size for *Nasuia*. If these differences translate into a smaller *N_e_*, such transmission bottleneck could further weaken purifying selection and explain the prevalence of nonsynonymous variants at intrahost scale in *Nasuia*. Over longer evolutionary timescales, fixation of variants affecting these remaining functions could contribute to functional erosion and help explain the history of relatively frequent replacement of betaproteobacterial endosymbionts across insects in the Auchenorrhyncha suborder (e.g., spittlebugs, cicadas, leafhoppers; McCutcheon and Moran 2007; McCutcheon and Moran 2010; Koga and Moran 2014; Łukasik et al. 2018).

Our results provide some window into understanding how selection acts alongside drift in *Karelsulcia* and *Nasuia*. At the genome scale, the overall low pN observed in both endosymbionts indicates that purifying selection remains operative, constraining amino-acid–altering mutations despite substantial mutational input (**Fig. 4**). In addition, the overlap between intrahost mutational hotspots and founder–contemporary substitutions suggests that short-term variation can feed into longer-term evolutionary change, raising the possibility that some mutations experience non-random persistence over time (**Fig. 5**). Although the mode of selection operates at a different hierarchical level than the intrahost variation examined here, host-level selection on endosymbiont haplotypes has been demonstrated in the aphid–Buchnera system (Perreau et al. 2021). Disentangling the relative contributions of selection and drift to the ultimate fate of the intrahost variants in our system will therefore require comparable multigenerational, host fitness data similar to the previous aphid-*Buchnera* study.

Our results further point to a process in which substantial mutational input is absorbed with modest impact on core functions. On short evolutionary and ecological time frames, extreme polyploidy may explain how high mutational input has limited functional disruption. Both *Karelsulcia* and *Nasuia* carry hundreds of genome copies per cell (Woyke et al. 2010). Such redundancy could allow new mutations to persist across some number of genomic copies, cells, and bacteriocyte-localized populations without immediately disrupting essential functions. Polyploidy has also been proposed to facilitate conventional DNA repair by providing intact templates for double-strand break repair. However, *Karelsulcia* and *Nasuia* lack key factors (e.g., *recA*), making such repair-based buffering unlikely in this system (Bennett and Moran 2013).The evolutionary role of ploidy in these bacterial symbionts remains essentially uncharacterized. Clarifying its extent, dynamics, and consequences will require single-cell or haplotype-resolved approaches that can distinguish mutations among genome copies and among cells.

The mitochondria offer a well-studied internal comparative system to understand basic hypotheses of the processes that shape intrahost variation in vertically transmitted endosymbionts.

Correlations between mitochondrial and endosymbiont evolutionary rates have been reported in some insect–symbiont systems at macroevolutioanry scales (Degnan et al. 2004; Arab et al. 2020; Arab and Lo 2021; Degnan et al. 2025). Mitochondria harbor higher levels of heteroplasmy, as expected for a genome that evolves faster than endosymbionts (Vasquez and Bennett 2022; Degnan et al. 2025). However, mitochondrial heteroplasmy did not covary with *Karelsulcia* and *Nasuia* nucleotide diversity across hosts, consistent with the decoupled substitution rates reported for related Hawaiian leafhoppers in the genus *Nesophrosyne* (Vasquez and Bennett 2022). Several biological differences likely limit mitochondria as a direct mechanistic analogy in our system. A well-characterized germline bottleneck during oogenesis reshapes heteroplasmy in ways that may not have clear analogs in endosymbiont transmission (Kobiałka et al. 2016; Jia et al. 2017; Klucnika and Ma 2019; Zaidi et al. 2019; Mao et al. 2020). Moreover, mitochondrial and endosymbiont genomes each retain very small sets of functionally specialized genes for oxidative phosphorylation versus nutrient biosynthesis and other cellular functions, respectively. As a result, selection can act independently on these metabolic modules, differentially shaping their genetic diversity. In our system, while mitochondria provide an internal benchmark confirming that intrahost variation is not unique to endosymbiont genomes, it further highlights distinct differences in cellular context, developmental bottlenecks, and gene content. It provides limited value as a direct mechanistic analogy for understanding the evolution of *Karelsulcia* and *Nasuia* and likely other insect endosymbionts. It will be important in those cases to determine whether mutational regimes and magnitudes are themselves coupled, or only substitution rates.

In summary, results from this study show that our intrahost genetic diversity framework can act as a *de facto* experimental-evolution platform for obligate endosymbionts. Non-culturability, host integration, and longer generation times impose logistical challenges greater than classic evolution experiments with free-living bacteria (e.g., *E. coli*; Barrick et al. 2009). Even so, the combination of high mutational input and predictable functional impact provides a baseline expectation for how endosymbiont genomes might respond to sustained or manipulated conditions. We anticipate that this framework will be broadly applicable to the diversity of other unculturable, vertically transmitted endosymbionts with highly reduced genomes. In such systems, quantifying intrahost diversity will help anchor comparative inferences about how short-term mutational and microevolutionary processes scale to shape long-term genome erosion.

From a broader biological and evolutionary perspective, our study shows that laboratory empirical systems can serve as exceptionally high-resolution natural experiments to address questions of endosymbiont evolution. The biases in mutation rate, functional impact, and locus-specific instability that sculpt endosymbiont genomes over macroevolutionary timescales are already evident in the segregating and newly fixed mutations we observe within hosts over only ∼11 years. By resolving how new mutations arise, which genes they affect, and how they persist or move toward fixation, we directly observe the mechanistic pathways through which intrahost processes translate into long-term genome erosion. Rather than distorting endosymbiont evolution, long-term host colonies expose, in real time, the processes that continually generate and shape diversity in these ostensibly “clonal” intracellular partners.

## Materials and Methods

### Insect model and DNA extraction

*Macrosteles quadrilineatus* were obtained from previously established lab-reared colonies (Bennett and Moran 2013). Colonies were maintained on barley plants (*Hordeum vulgare*; aged one to four weeks) with a 12L:12D photoperiod (Percival Scientific, Inc., model I-36VL). Plants were grown from 1.5 g of seed in Miracle-Gro potting mix fertilized with Osmocote Smart-Release fertilizer.

We collected 20 individual adult *M. quadrilineatus* from two experimental cohorts (metadata available under **Supplementary Table 1**). One cohort (5 females, 5 males) was exposed to 42 °C for 24 h, and the other cohort (5 females, 5 males) was maintained at 25 °C. All individuals were flash-frozen with liquid nitrogen and processed identically for DNA extraction.

Total genomic DNA was extracted using the Quick-DNA/RNA Microprep Plus Kit following the manufacturer’s Tough-to-Lyse Samples protocol (Zymo Research, Irvine, CA, USA). DNA quality and concentration were measured via Nanodrop and Qubit 4.0 Fluorometer (Invitrogen, Carlsbad, CA, USA).

### Library preparation and sequencing

The 20 individual libraries were generated in two sequencing batches (n = 10 per batch; **Supplementary Table 1**). Barcode-indexed libraries were generated using Watchmaker DNA Library Prep Kit with Fragmentation following the manufacturer’s recommendations (Watchmaker Genomics, CO, USA). Libraries were checked with a Bioanalyzer 2100 (Agilent, Santa Clara, CA), quantified with a Qubit 4.0 Fluorometer (Invitrogen, Carlsbad, CA, USA), and pooled at equimolar ratios. The pools were quantified by qPCR with a Kapa Library Quant kit (Kapa Biosystems-Roche).

All libraries were sequenced as paired-end 150 bp read lengths with an average insert size of ∼320bp (Picard CollectInsertSizeMetrics; duplicates excluded; Broad Institute 2025). Libraries from batch 1 were first sequenced on an AVITI platform (Element Biosciences, San Diego, CA) across two runs that were merged to achieve the desired depth. Libraries from batch 2 were sequenced on a NovaSeq X Plus (Illumina, San Diego, CA) to take advantage of the platform’s higher throughput. To obtain platform-independent technical replication, all 20 libraries were additionally sequenced a second, independent NovaSeq X Plus lane. We used this replicate dataset to validate variant detection in downstream analyses. Library preparation and sequencing were performed at the DNA Technologies Core Facility at the University of California, Davis.

### Read pre-processing and binning

Raw reads were quality-checked with FASTQC v0.11.9 (Andrews 2010). Low-quality reads and adapter sequences were removed using Trimmomatic v0.39 (Bolger et al. 2014). We separated host and bacterial reads using BBSplit (Bushnell 2014) by binning reads to reference genomes of *M. quadrilineatus* (NCBI RefSeq: GCF_028750875.1), *Karelsulcia* (GenBank: CP006060.1), and *Nasuia* (GenBank: CP006059.1). Because the host reference genome already includes the mitochondrial chromosome (GenBank: NC_034781.1), mitochondrial reads were included in the host read pool and subsequently extracted using PrintReads and RevertSam in Picard v.2.26.2 (Broad Institute 2025).

### *De novo* population consensus genome assembly

To minimize reference bias in downstream variant calling and genetic diversity analyses, we constructed contemporary population consensus genomes for *Karelsulcia*, *Nasuia,* and mitochondria. Previous publicly available references were generated from field-collected samples in 2012 and may not reflect the genetic composition of our sampled population. To generate contemporary references, we first merged reads assigned to each bacterial symbiont and mitochondria from all individuals and both sequencing runs. Because pooled depths were extremely high (with *Nasuia* having the lowest pooled coverage at ∼3,000x), we normalized the coverage to 100x with BBNorm (Bushnell 2014) to produce an even k-mer profile and prevent de Brujin graph inflation during assembly. Normalized read sets for *Karelsulcia* and *Nasuia* were assembled using SPAdes v4.2.0 (Prjibelski et al. 2020). Mitochondrial reads were assembled with GetOrganelle v1.7.7.1 (Jin et al. 2020).

For each initial assembly, we identified the largest scaffolds that matched expected genome characteristics (e.g., length, GC content, and coverage). We evaluated synteny against published reference genomes using Mauve (Darling et al. 2004). Assemblies were oriented to match public references and inspected for local mis-assemblies and sequence errors using Pilon v1.24 (Walker et al. 2014). This effort confirmed that no additional corrections were required. Final high-quality assemblies were annotated in Geneious Prime 2025.2.1 (https://www.geneious.com) using custom reference-derived databases (95% similarity threshold). Percent pairwise nucleotide identity between each population consensus genome and its published reference was estimated from global pairwise alignments in Geneious Prime 2025.2.1 (https://www.geneious.com). For the mitochondrial genome, the hypervariable A+T-rich control region (positions 14,486-16,626) was excluded from the alignment.

To improve variant detection near circular junctions, as is recommended for bacteria and mitochondria (Van der Auwera & O’Connor, Brian D, 2020; see **Variant discovery and annotation**), we also generated a shifted version of each population consensus genome by rotating the sequence by half its length using ShiftFasta in GATK v4.2.6 (McKenna et al. 2010).

### Variant discovery and annotation

We followed GATK’s Best Practices for variant calling in microbes (github.com/broadinstitute/GATK-for-Microbes) and mitochondria (github.com/gatk-workflows/gatk4-mitochondria-pipeline). To perform variant calling, we aligned binned reads to their respective population consensus genomes, in both the original and shifted forms, using Bowtie2 v1.3.0 (Langmead and Salzberg 2012) in local alignment mode. We also generated technical-duplicate alignments from each sequencing run. The alignments were converted and sorted using SAMtools v1.13 (Li et al. 2009). PCR duplicates were marked using MarkDuplicates in Picard v.2.26.2 (Broad Institute 2025).

To avoid run-specific coverage biases in downstream variant discovery, we first estimated average coverage for each BAM using SAMtools v1.13 (Li et al. 2009). We then down-sampled the higher-depth BAM to match the lower-depth BAM using DownsampleSam in Picard v.2.26.2 (Broad Institute 2025). This procedure produced four normalized BAM files for each endosymbiont or mitochondria in host individuals: (i.) Run 1-original, (ii.) Run 1-shifted, (iii.) Run 2-original, and (iv.) Run 2-shifted. Variants were identified independently from each of the four normalized BAM files using Mutect2 in GATK v4.2.6 (McKenna et al. 2010).

To obtain a unified variant dataset for each sequencing run, we converted variant positions from the shifted-reference Variant Call Format (VCF) files back into the original-reference coordinate system and then merged these with the corresponding original-reference callsets using Picard v.2.26.2 (Broad Institute 2025). Shifted-reference callsets were identical to unshifted ones, confirming that variant detection across the circular junction was fully resolved. Final unified VCFs were deduplicated, multiallelic variants were decomposed and split, and indels were left-aligned using bcftools v1.14 (Danecek et al. 2021).

We annotated each unified VCFs using SnpEff v5.2 (Cingolani et al. 2012) with custom databases constructed from the population consensus genomes. The resulting annotated callsets (Run 1-unified and Run 2-unified) were then carried forward to downstream validation, where they were subsequently reconciled (see below).

### Variant validation

Both NovaSeq X Plus and AVITI short-read sequencing platforms exhibit very low baseline substitution error rates, yielding 85% of bases in the highest quality bin (Q40, less than one error per 10, 000 bases; Arslan et al. 2024; Yao 2025). However, the extreme A+T bias and frequent homopolymers in endosymbiont and mitochondria genomes are known to increase sequencing artifacts (Stoler and Nekrutenko 2021). To validate the detection of true segregating variants from sequencing errors, we implemented several complementary quality-control steps independently for each sequencing run. We then compared the resulting callsets across our deeply sequenced technical duplicates.

We first applied an allele-frequency (AF) filter threshold of 1-98% to each independent sequencing data set, following (Wei et al. 2019). Because each library represents a pooled population of endosymbiont and mitochondria genomes, the variant allele fraction reported by Mutect2 corresponds to the population AF. The 1% lower bound is substantially higher than error rates for short-read sequencing platforms. This approach conservatively removes rare variants likely driven by technical noise rather than biological signal. The 98% upper bound excludes near-fixed alleles, which provide little information about segregating variation.

We further filtered variant calls using FilterMutectCalls GATK v4.2.6 (McKenna et al. 2010).

Depending on the data source, callsets were processed in microbial or mitochondrial mode. This filtering step applies a probabilistic model to remove false positives arising from strand bias, low mapping or base quality, homopolymer-associated errors, and other alignment artifacts. FilterMutectCalls annotates each site that passes all quality checks with a ‘PASS’ flag. Variants with any non-PASS flag, were discarded from downstream analyses.

Finally, as noted above, we intersected the two unified callsets (Run 1-unified and Run 2-unified) using bcftools v1.14 (Danecek et al. 2021). Technical-replicate concordance is widely recognized as one of the most robust strategies for validating variants, particularly in error-prone or compositionally biased genomic contexts (Robasky et al. 2014; McCrone and Lauring 2016; Kim et al. 2019; Wei et al. 2019). We therefore retained variants only if they were independently recovered in both sequencing runs and passed all upstream QC steps under identical calling and filtering conditions.

### Assessment of batch and treatment effects on variant profiles

To test whether library batch, sample cohort (25 °C vs 42 °C), or host sex introduced systematic structure in variant profiles, we analyzed the presence–absence of segregating sites across samples. For each genome (*Karelsulcia*, *Nasuia*, mitochondria), we constructed a binary matrix with samples and validated variant positions, where each entry indicated whether that variant was detected in that sample.

From this matrix, we computed Jaccard distances among samples and used PERMANOVA (adonis2 in the R Vegan package) to partition variance in mutational composition for cohort, library batch, and sex for each endosymbiont and mitochondria. PERMANOVA identified modest batch-associated effects for two endosymbionts (R^2^ = ∼0.10, nominal P < 0.05 for both), but no effects of cohort or host sex (**Supplementary Table 4**).

To test whether this signal was driven by specific variants, we applied Fisher’s exact tests at each position to identify variants that were disproportionately present in one batch. We did not detect any batch-associated sites. Because sparse, rare variants can nonetheless generate spurious multivariate structure, we further asked whether the PERMANOVA batch signal could arise by chance. We generated a null distribution of PERMANOVA P-values by shuffling batch labels (1,000 permutations) while keeping the distance matrix fixed. The observed batch P-value fell well within this null distribution, indicating that the nominal batch-associated signal is consistent with chance rather than a genuine batch effect (**Supplementary Fig.2)**.

### Variant class analyses

To compare overall level of intrahost variation, we first calculated variant densities by normalizing total variant counts to callable genome size per kilobase (count / callable bases x 1,000). We obtained counts for three variant classes: (i) single nucleotide variants (SNVs), (ii.) multiple nucleotide variants (MNVs), and (iii.) small insertions and deletions (indels; < 50 bp) using bcftools v1.14 (Danecek et al. 2021). Callable genome sizes, defined as the total number of high-coverage, non-masked bases, were extracted from the Mutect2 statistics output. We compared total and class-specific variant densities between *Karelsulcia* and *Nasuia* using Wilcoxon signed-rank tests. Correlations with mitochondria variant class profiles were tested with Spearman’s rank correlations.

We quantified single-base mutation rates and transition/transversion (Ti/Tv) ratios. As in the variant density analysis, SNV counts were normalized by the corresponding callable positions for each of the six-directional base-change classes. Callable positions for each class were computed using custom scripts. Raw counts for all six classes were obtained from SnpEff-generated summary reports. We compared differences in SNV rates using Kruskal-Wallis, followed by Dunn’s post hoc test. Ti/Tv ratios were then aggregated from these normalized SNV rates and compared using Wilcoxon signed-rank tests.

We also characterized the distribution of indel lengths and the fraction of indels occurring in repeat-sequence regions. We extracted indel lengths and genomic coordinates from each VCF using custom scripts. We compared differences in indel-length distributions using Fisher’s exact test. We then identified homopolymer and short tandem repeat (STR) tracts in each population consensus genome using MISA v2.1 (Thiel et al. 2003; Beier et al. 2017). We defined repeat sequences as homopolymers and STRs with a track length ≥ of 4 bp. Indel coordinates were then intersected with these tracts using GenomicRanges (Lawrence et al. 2013). Indel densities for each repeat category were calculated by normalizing indel counts to the total genomic sequence length of the corresponding sequence class.

### Genetic diversity metrics

We measured genetic diversity within endosymbiont and mitochondria populations using two AF-based metrics: (i) the fraction of intermediate-frequency variants, and (ii) nucleotide diversity (π). To improve accuracy, we re-estimated AFs for all validated variants from merged sequencing duplicates. We defined intermediate-frequency variants as those with 0.25 ≤ AF ≤ 0.75 (Russell and Cavanaugh 2017; Kelly and Hughes 2019; Mérel et al. 2021). Nucleotide diversity in our study is equivalent to expected heterozygosity at the nucleotide level following (Nei and Kumar 2000). We calculated π as: π = 1 - ∑i=1^k^ (p_i_)^2^, where p*_i_* is the frequency of the i^th^ allele at a site, and k is the number of alleles (Nei 1973). To calculate genome-wide and local diversity, we averaged π across all sites (π = 0 at invariant sites) and 1 kb non-overlapping windows along each population consensus genome. To measure how local diversity is spatially distributed, we quantified heterogeneity of window π using Gini coefficients and Shannon diversity indices (Tomezsko et al. 2020; Fuhrmann et al. 2021). We also evaluated spatial autocorrelation of diversity using Moran’s I (Moran 1948; Feltus et al. 2004).

### Functional analyses

To analyze functional impacts of SNVs, we estimated rates of nonsynonymous (pN) and synonymous single-base changes (pS) using a codon-aware AF-weighted framework following Shenhav and Zeevi 2020 as implemented by Kiefl et al. 2023. For each codon in the population consensus genomes, we tabulated all possible single-base changes (three positions x three alternative bases) and counted how many of these potential changes would be synonymous (n_S_) versus nonsynonymous (n_n_). We then mapped each observed variant to its corresponding codon and assigned each variant’s functional class (nonsynonymous or synonymous) using SnpEff annotations. For each coding site, we calculated pS or pN by multiplying the variant AF by 1⁄n_S_ or 1⁄n_n_, respectively. Gene-level pN and pS were obtained by summing these site-wise rates across all coding positions in the relevant regions.

Because an analogous codon-opportunity model is not available for indels, we defined a conceptually comparable AF-weighed coding indel rate at the gene level. For each gene in each endosymbiont population, we summed the AFs of all validated indels within its coding sequence. Similar to Waneka et al. 2021, we then normalized this sum by the gene’s coding length in nucleotides.

To evaluate whether particular biological processes exhibit distinct variation patterns, we assigned functional categories by mapping all protein-coding genes to Clusters of Orthologous Groups (COG) with eggNOG-mapper v2 (Cantalapiedra et al. 2021). Genes lacking definitive COG annotations were manually curated based on previous endosymbiont genome studies (Bennett et al. 2014). We compared pN, pS, and indel rate across COG functional categories using Kruskal-Wallis, followed by Dunn’s post hoc tests.

### Fonder-contemporary divergence

Although the population consensus genomes are highly similar to the original founder references, we observed several substitutions and indels between them. To quantify how these changes are distributed across current endosymbiont populations, we re-ran the variant calling and annotation pipeline described above, mapping reads to the original founder references for *Karelsulcia* (GenBank: CP006060.1) and *Nasuia* (GenBank: CP006059.1). For this analysis, we applied the same quality and duplicate-reproducibility filters but did not impose the AF ≤ 0.98 upper bound, to capture substitutions relative to the founder state. We then cataloged sites where a contemporary allele constituted the major allele (AF > 0.75) in at least one endosymbiont population. For these fixed changes, we recorded their functional effect class using SnpEff annotations.

## Data Availability

All raw sequencing reads are deposited at NCBI under BioProject PRJNA1391824.

## Code Availability

Custom scripts used for data processing and analyses are available at https://github.com/younghwankwak/endosymbiont-intrahost-diversity

## Acknowledgements

We thank Dr. Nancy Moran, Dr. Martin Kaltenpoth, and Dr. Kaden Muffett for insightful comments and constructive suggestions on the early manuscript draft. We thank Toxtli Huitzilopochtli for assistance with insect rearing and sample collection. This work was supported by the National Science Foundation (NSF) under Grant No. DBI-2214038. Library preparation and sequencing were carried out at the DNA Technologies and Expression Analysis Cores of the University of California, Davis Genome Center, supported by NIH Shared Instrumentation Grant No. 1S10OD010786-01. Computational resources were provided in part by CENVAL-ARC (NSF Grant No. 2346744) and by the Pinnacles cluster at the Cyberinfrastructure and Research Technologies (CIRT) at the University of California, Merced (NSF MRI Grant No. ACI-2019144). An AI-based language tool was used for limited assistance with code debugging and language editing; all analyses, interpretations, and conclusions were performed and verified by the authors.

## Author contributions

We followed CRediT (Contributor Roles Taxonomy) guidelines. **Y.K.:** Conceptualization; Methodology; Software; Validation; Formal Analysis; Investigation; Data Curation, Writing – Original Draft; Writing – Review & Editing; Visualization; Project Administration. **G.M.B.:** Conceptualization; Resources; Writing – Review & Editing; Supervision; Funding Acquisition.

