## Supplementary Figures 1-3 for "Intrahost mutational dynamics parallel long-term genome evolution in endosymbionts"

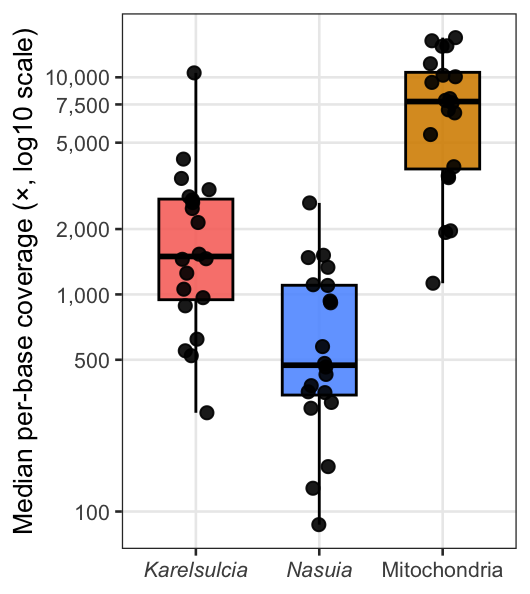

**Supplementary Figure 1. Sequencing coverage of symbionts and mitochondria.** Median per-base coverage for *Karelsulcia*, *Nasuia*, and mitochondria across 20 populations. Coverage is shown on a log_10_ scale.

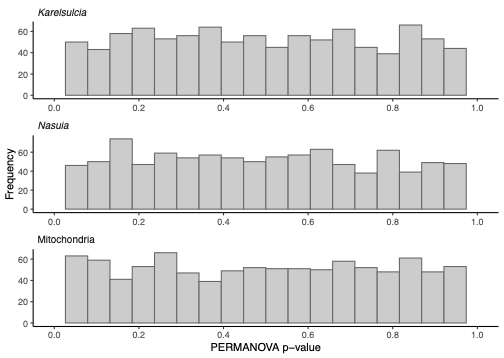

**Supplementary Figure 2. Null distribution of PERMANOVA p-values for batch effects.** Distribution of PERMANOVA p-values were computed from pairwise genetic distance matrices derived from variant presence-absence data under 1,000 permutations. P-values do not deviate from the null distribution, providing no evidence for batch-associated structure.

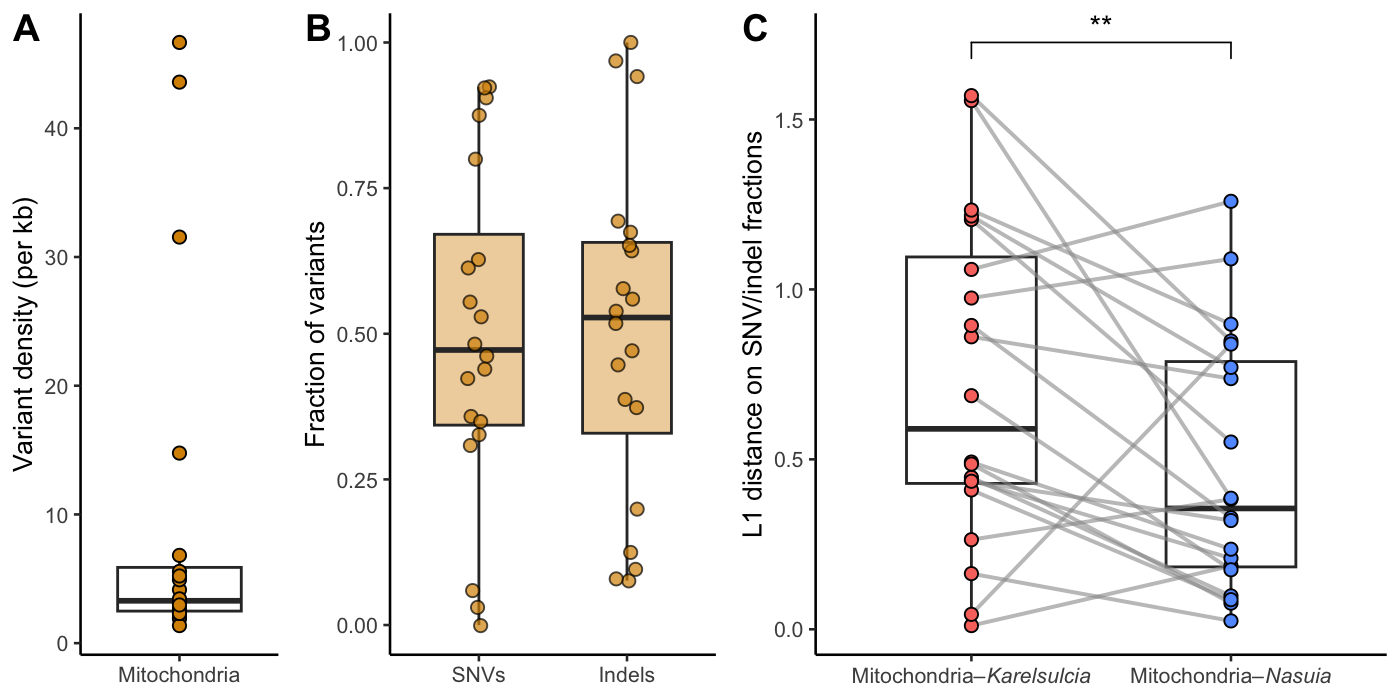

**Supplementary Figure 3. Mitochondrial variant density and class composition across 20 populations. (A)** Variant density of mitochondrial genomes expressed as variants per kilobase. **(B)** Fractional composition of mitochondrial variants, partitioned into single-nucleotide variants (SNVs), and insertions and deletions (indels). **(C)** L1 distance between mitochondrial and endosymbiont variant class composition (SNV/indel fractions). Mitochondrial variant profiles are more similar to *Nasuia* than to *Karelsulcia* (**, *P* < 0.01; Wilcoxon signed-rank test).

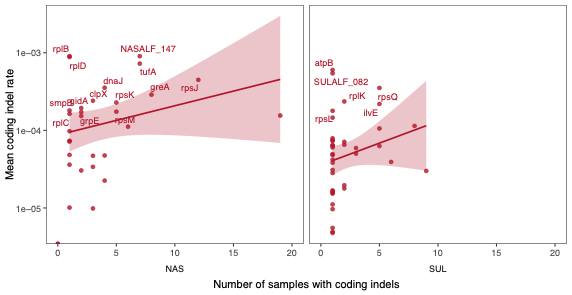

**Supplementary Figure 4. Coding-sequence indel rates in endosymbionts** Mean coding indel rates per gene are plotted against the number of populations in which coding indels were detected. Each point represents a gene. Rates are shown on a log scale to illustrate variation across genes and populations. NAS, *Nasuia*; SUL, *Karelsulcia*.
